# ADAM10 tailors extracellular vesicles for content transfer rather than signaling by contact

**DOI:** 10.64898/2026.02.12.705562

**Authors:** R. Ghossoub, L. Goullieux, S. Audebert, L. Hyka, E. Jaafar, S. Granjeaud, M. Methia, S. Thuault, R. Leblanc, G. David, P. Zimmermann

## Abstract

Etxtracellular vesicles (EVs) support cell-to-cell communication, both in physiological and pathological contexts and emerge as new biomarkers and potential therapeutics. Yet, despite EVs holding huge promises, our understanding of core mechanisms governing EV signaling remains significantly underdeveloped.

Our previous work indicated that syndecans and their cytosolic adaptor syntenin control the biogenesis of a major subset of small EVs (sEVs). Here we show that syndecans control the accumulation of ADAM10 into sEVs. ADAM10 promotes the formation of sEVs enriched in cleaved receptors (reducing sEV corona), supports the sorting of proteins with intracellular functions, and tailors sEVs for the delivery of their internal content, e.g. syntenin, into the cytosol of recipient cells. Conversely, inhibition of ADAM10 favors the production of sEVs bearing full-length, signaling-competent receptors/ligands and enhances contact-dependent signaling. These findings uncover a protease-regulated switch that tailors sEV composition and signaling modality, providing important new mechanistic insights into the core molecular pathways supporting EV-mediated communication.

## Introduction

Extracellular vesicles (EVs) play a crucial role in intercellular communication by transporting proteins, lipids, and nucleic acids between cells. They are involved in a wide range of physiological and pathological processes, including immune regulation, tissue repair, and cancer progression. Despite growing interests in EVs for diagnostics and therapeutics, core molecular mechanisms governing EV exchanges and modes of signaling remain poorly understood.

EVs support intercellular communication through multiple signaling modalities. One key mechanism involves direct interaction between EV surface molecules and receptors on target cells, triggering signaling pathways, probably without the necessity for vesicle internalization. This receptor-mediated contact-dependent signaling enables rapid cellular responses and modulation of downstream effectors (Yáñez-Mó *et al*, 2015; Jahnke & Staufer, 2024). Alternatively, EVs can fuse with contacting membranes, directly at the cell surface or (more likely) following internalization via various endocytic routes, including phagocytosis and macropinocytosis, leading to the delivery of EV cargo (comprising proteins, lipids, and various RNA species) to the membranes and the cytoplasm of recipient cells. The transfer of these bioactive molecules can induce significant phenotypic changes in recipient cells by influencing gene expression and cellular metabolism (van Niel *et al*, 2018, 2022). Nevertheless, whether and how EV embedded or inner cargo can be efficiently delivered into the cytosol of EV recipient cells remains debated (Ngo *et al*, 2025). Also, mechanisms that may control EV surface composition or corona are under-investigated (Försönits *et al*, 2025; Buzas, 2022).

Our previous work identified the PDZ protein syntenin and its major binding partners, the syndecans (SDCs), as key drivers of small EV (sEV) biogenesis. In particular, we showed that syntenin promotes intraluminal vesicle (ILV) formation at endosomal membranes by adapting endocytosed SDC to ALIX, a process mechanistically reminiscent of viral budding (Baietti *et al*, 2012). Notably, the SDC-syntenin pathway can contribute to up to 50% of the sEV production (Baietti *et al*, 2012; Ghossoub *et al*, 2014; Imjeti *et al*, 2017). Along this biogenesis pathway, SDCs undergo proteolytic cleavage, resulting in the enrichment of a membrane-spanning C-terminal fragment of SDC (SDC-CTF) within sEVs (Baietti *et al*, 2012; Roucourt *et al*, 2015; Ghossoub *et al*, 2022).

In a recent study (Castro-Cruz *et al*, 2023), we found that SDC-CTF levels on sEVs correlate well with sEV abundance and may serve as an accurate molecular indicator of sEVs. However, the mechanisms regulating sEV-SDC cleavage, as well as the consequences of this process on EV-mediated signaling, remained largely unknown. Here, we aimed to identify proteases responsible for SDC cleavage and to elucidate their functional impact for sEV biology.

## Results

### ADAM10 (not ADAM17) supports the proteolytic processing of sEV-syndecans

To identify proteases potentially involved in sEV-related SDC cleavage, we first immunoprecipitated GFP-tagged SDC4 from transfectant MCF7 cell extracts and analyzed associated proteins by differential proteomics (comparing these IPs to GFP immunoprecipitates). Among SDCs expressed in MCF7 cells, SDC4 and SDC1 are the most abundant (Ghossoub *et al*, 2020), SDC4 being near ubiquitous (common to multiple different cell types). The sole protease peptides identified corresponded to ADAM10 and ADAM17, two metalloproteases implicated in the shedding of a plethora of membrane receptors (Huovila *et al*, 2005; Pruessmeyer & Ludwig, 2009) (Fig. 1A). Both interactions were validated in co-immunoprecipitation experiments (Fig. 1B).

**Figure 1.**
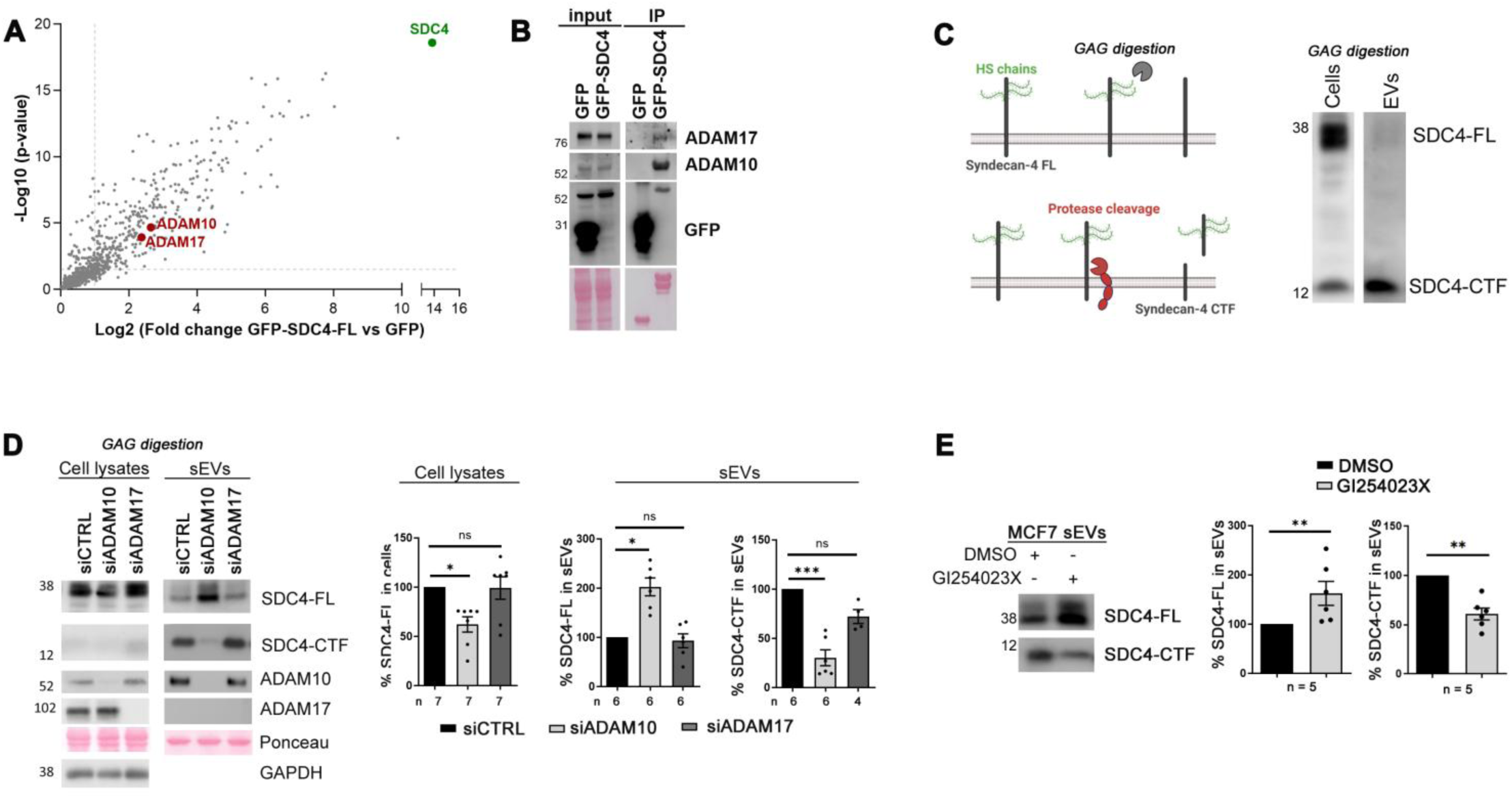
Both ADAM10 and ADAM17 interact with SDC4, but solely ADAM10 (not ADAM17) downregulation or inhibition modifies sEV-SDC4. **A.** Volcano plot illustrating the proteins associated with GFP-SDC4 according to differential proteomics, with log2 fold change (x-axis) and −log10 (p-value) (y-axis). Data were obtained by GFP-pulldown from total extracts of MCF7 cells overexpressing GFP-SDC4 compared to MCF7 cells overexpressing GFP alone. Data result from three different experiments processed three times. SDC4 (the bait) is represented in green, ADAM10 and ADAM17, the two metalloproteases associated with SDC4 are represented in red. The other proteins are represented in grey. The proteins showing non-significant differences are represented below the dotted lines (threshold at 1 for the x-axis, difference =2 and 1.5 for y-axis; p-value = 0.3). **B.** Co-immunoprecipitation experiments from MCF7 cell lysates, confirming the association of endogenous ADAM10 and ADAM17 with GFP-SDC4, but not GFP. The GFP precipitates (IP) were subjected to Western blot with antibodies recognizing proteins as indicated on the right. **C**. **Left:** Scheme illustrating the complexity of SDC4 structure and processing. (Upper part) SDC4 full-length (FL) core protein (black) is substituted with glycosaminoglycan chains (GAG) of the heparan sulfate type (green). The trimming of GAG chains by specific enzymes is necessary for the FL protein to migrate at a discrete band in SDS-PAGE and to be easily detectable by immunoblotting. (Lower part) SDC4 core protein can be cleaved by proteases generating two main fragments: an N-terminal fragment comprising most of the GAG-substituted extracellular domain (ECD) and a C-terminal fragment (CTF) comprising the remainder of the ECD, the membrane-spanning and the cytoplasmic domain. **Right:** Western blot illustrating the signals obtained, after GAG-digestion, for FL SDC4 and SDC4 CTF in the cells and the sEV enriched fraction obtained after differential ultracentrifugation of the conditioned extracellular media. Signals were obtained with an antibody recognizing the intracellular domain of SDC4. Note that the FL form of SDC4 (SDC4-FL) abounds in cell lysates, while the CTF is less abundant. On the contrary, the SDC4-CTF is abundant and the SDC4-FL is barely detectable in sEVs. **D.** MCF7 cells downregulated for ADAM10 (siADAM10) or ADAM17 (siADAM17) and control cells (siCTRL) were evaluated for SDC4 FL and CTF abundance in cells and sEVs by Western blot after GAG-digestion. Histograms represent the mean signal intensity for indicated proteins relative to the signal in control cells, ± SEM. Statistical analysis was performed using the Kruskal-Wallis one-way non-parametric ANOVA test (* P < 0.05, *** P < 0.001, n.s. non-significant). **E**. sEVs secreted by MCF7 cells inhibited for ADAM10 activity, (GI254023X) versus controls (DMSO) were isolated from conditioned media by differential ultracentrifugation. sEVs were analyzed by Western blot after GAG-digestion to evaluate the levels of SDC4 FL and CTF. Histograms represent the mean signal intensity for indicated proteins relative to the signal in control cells, ± SEM. ** P < 0.01 (unpaired non-parametric Mann–Whitney test).

As previously mentioned, SDCs undergo proteolytic cleavage and, compared to cell lysates, sEVs are substantially enriched in C-terminal fragment of SDC (SDC-CTF) (Fig. 1C, Fig. S1A). To determine the effects of ADAM10 or ADAM17 on the proteolytic processing of sEV-SDCs, we first downregulated ADAM10 and ADAM17 expression using siRNAs. Both siRNAs were efficient, yet, solely ADAM10 (not ADAM17) was detected in sEVs and influenced sEV SDC4 loading (Fig. 1D). sEVs secreted by cells where ADAM10 is downregulated, contain two-fold more SDC4 that is full-length and four-fold less SDC4 CTF. This result is not unique for SDC4, as the cleavage of SDC1 and levels of SDC1 ICD accumulating in sEVs appear similarly regulated by ADAM10 (Fig. S1B). Of note, SDC-FL requires digestion of its GAG chains for detection in western blot (as shown in Fig. S1C). As can be calculated from the combination of the data obtained in multiple independent experiments the total levels of SDC intracellular domain (ICD) accumulating in sEVs, present in either full-length or CTF forms of SDC, remain nearly unchanged (Fig. S1D-E).

Interestingly, these effects of ADAM10 appear to strictly depend on its enzymatic activity, as similar results were obtained upon treatment of cells with the ADAM10 inhibitor GI254023X (Fig. 1E for SDC4, Fig. S1F for SDC1). Moreover, the effects of ADAM10 knock-down were reproduced in different ADAM10-KO cells, including MCF7 (Fig. S2A), MDA-MB-468 (Fig. S2B) and SKBR3 (Fig. S2C) indicating that ADAM10 effects are not restricted to one cellular model or short-term downregulation/inhibition. Of note, ADAM10 KO had no major influence on the number of secreted particles (Fig. S2 D-F). Altogether these data identified ADAM10 as key regulator of sEV-SDC processing.

### Syndecans control ADAM10 sorting into sEVs, whereas ADAM10 does not affect the loading of sEVs with syntenin, ALIX and tetraspanin

Having identified ADAM10 as regulator of sEV SDC-processing, we next investigated whether SDC, in turn, may influence the incorporation of ADAM10 into sEVs. Using siRNA targeting SDCs, we observed that the downregulation of SDCs is accompanied by a decrease of ADAM10 levels in sEVs (Fig. 2A siSDC4, Fig. S3A siSDC1). As previously shown (Baietti *et al*, 2012), downregulation of SDC1 or SDC4 does not affect the cellular levels of syntenin, ALIX, CD81 and CD9, but impacts sEV compositions for these markers (Fig. 2A; Fig. S3A). Interestingly, downregulation of SDC1 or SDC4 was accompanied by a significant, near two-fold increase of cellular ADAM10 (Fig. 2A; Fig. S3A). Microcopy experiments indicated that (upon SDC-downregulation) ADAM10 accumulates inside CD63-positive cellular structures (Fig. 2B; Fig. S3C). Moreover, on the contrary to SDC downregulation, ADAM10 downregulation had no impact on the loading of syntenin, ALIX, and tetraspanin (CD63, CD81 and CD9) into sEVs (100K pellet, Fig. 2C). Persistence of SDC-dependent cargo would appear consistent with unchanged total levels of SDC intracellular domain (ICD) in sEVs (Fig. S1 D-E). Of note, ADAM10 or SDC silencing did not affect sEV size and number in nanoparticle tracking analysis (NTA) (Fig. 2D, S3B) or microfluidic resistive pulse sensing (MRPS) measurements (Fig. S2D-F). Similar results were obtained upon ADAM10 enzymatic inhibition (Fig. S3D). Collectively, these observations indicate that SDCs control the sorting of ADAM10 in sEVs and that ADAM10 affects the form (FL versus CTF) of SDC that is accumulating, but has no impact on the SDC-associated sEV markers implicated in sEV biogenesis (syntenin, Alix and tetraspanins).

**Figure 2.**
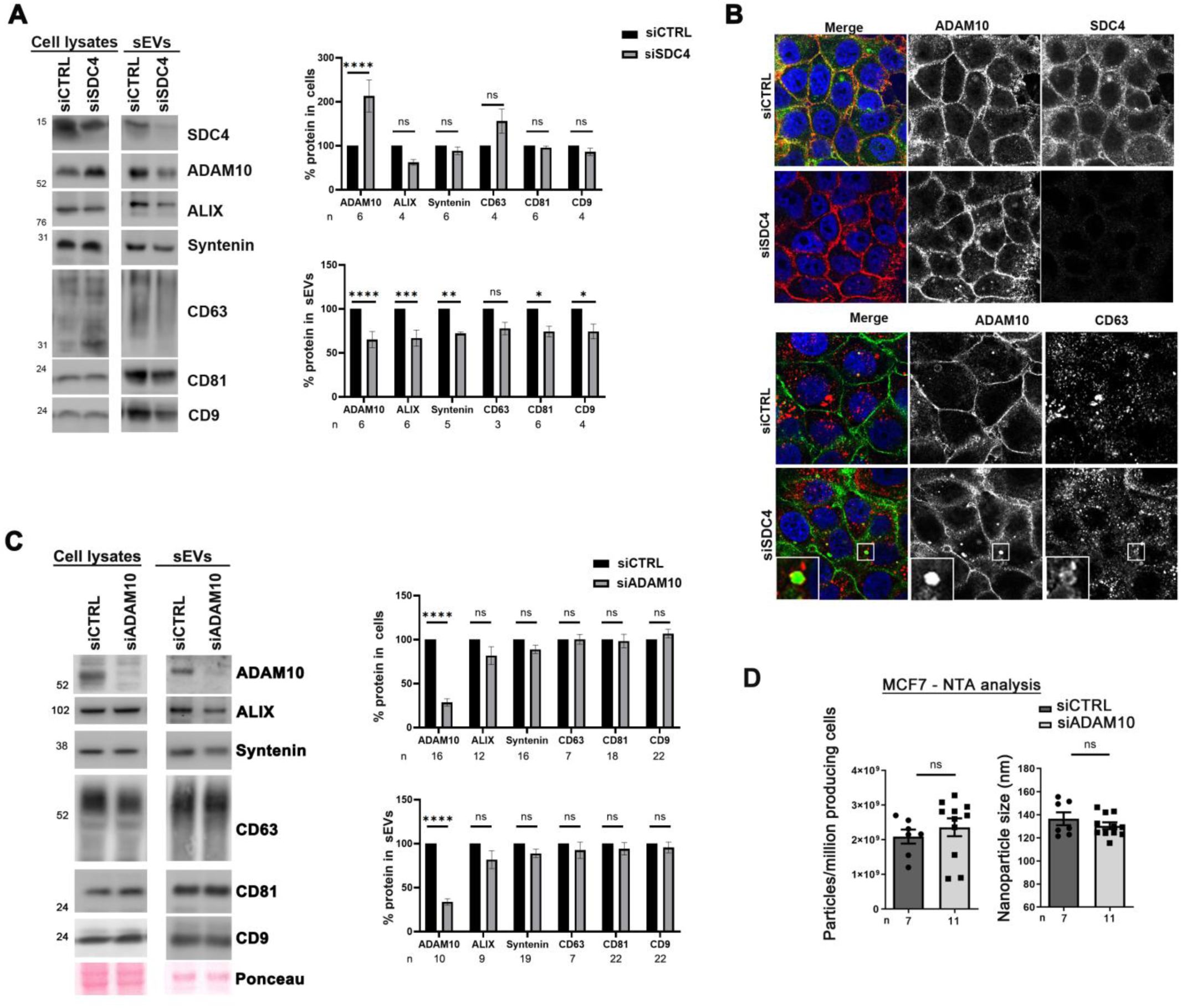
ADAM10 loading in sEVs is SDC4-dependent, yet ADAM10 downregulation does not change syntenin, ALIX and tetraspanins (CD9, CD63, CD81) levels in sEVs. **A.** sEVs secreted by SDC4-downregulated cells (siSDC4) versus controls (siCTRL) were isolated by differential ultracentrifugation. sEVs were further analyzed by Western blot, testing for several markers, as indicated. Histograms represent mean signal intensities relative to signals obtained for controls, ± SEM. * P < 0.05, ** P < 0.01, *** P < 0.005, **** P < 0.001, n.s. non-significant (two-way ANOVA). **B.** (Upper part) Representative confocal micrographs showing the steady-state distribution of endogenous ADAM10 (red in merge) in MCF7 cells transfected with control (siCTRL) or siRNA targeting SDC4 (siSDC4). SDC4 was stained with antibody recognizing the intracellular domain of SDC4 (green in merge). (Lower part) Representative confocal micrographs showing the steady-state distribution of endogenous ADAM10 (green in merge) together with CD63 (red in merge) in MCF7 cells transfected with control (siCTRL) or siRNA targeting SDC4 (siSDC4). In merge, nuclei are stained with DAPI (blue). **C.** sEVs secreted by ADAM10-downregulated cells (siADAM10) versus controls (siCTRL) were isolated by differential ultracentrifugation. Cell lysates from parental cells or sEVs were further analyzed by Western blot, testing for several markers, as indicated. Note that ADAM10 downregulation does not impact the loading of Alix, syntenin, and the tetraspanins CD63, CD81 and CD9. Histograms represent mean signal intensities in sEVs relative to signals obtained for controls, ± SEM. * P < 0.05, ** P < 0.01, *** P < 0.001, n.s. non-significant (two-way ANOVA). The value ‘n’ indicates the number of independent experiments. **D.** sEVs secreted by MCF7 cells downregulated for ADAM10 (siADAM10) or control cells (siCTRL) were isolated by differential ultracentrifugation from conditioned media and further analyzed by Nanosight (NTA) to determine concentration and size, as indicated. n indicates the number of independent experiments; each point represents one independent experiment.

### ADAM10 downregulation changes sEV composition

To further clarify the impact of ADAM10 on sEV composition, we performed label free mass-spectrometry analysis of sEVs secreted by MCF7 ADAM10-KO cells, comparing these to sEVs secreted by WT cells. We observed that, out of the 5325 proteins identified in sEVs, 426 proteins were significantly upregulated in ADAM10-KO sEVs, whereas 608 proteins were downregulated. Thus, ADAM10 affects approximatively 20% of the proteins composing sEV (Fig. 3A and S4A, Supplementary table 1). Proteomic profiling of MCF7 WT and ADAM10-KO cells identified a total of 7136 proteins, with 204 proteins and 196 proteins respectively upregulated and downregulated in ADAM10-KO cells. Thus, ADAM10 affects approximatively 5% of the cellular proteome (Fig. S4A, Supplementary table 2). Functional annotation revealed that cytosolic regulatory proteins including translational and mitochondrial proteins are enriched in EVs secreted by WT cells, while several proteins involved in cell-cell and cell-extracellular matrix communication are more abundant in EVs from ADAM10-KO cells (Fig. 3B). Overall these results highlight that ADAM10 has more impact on the proteome content of sEVs than on the cellular proteome and point to a differential packaging of sEV cargos under ADAM10 regulation.

**Figure 3.**
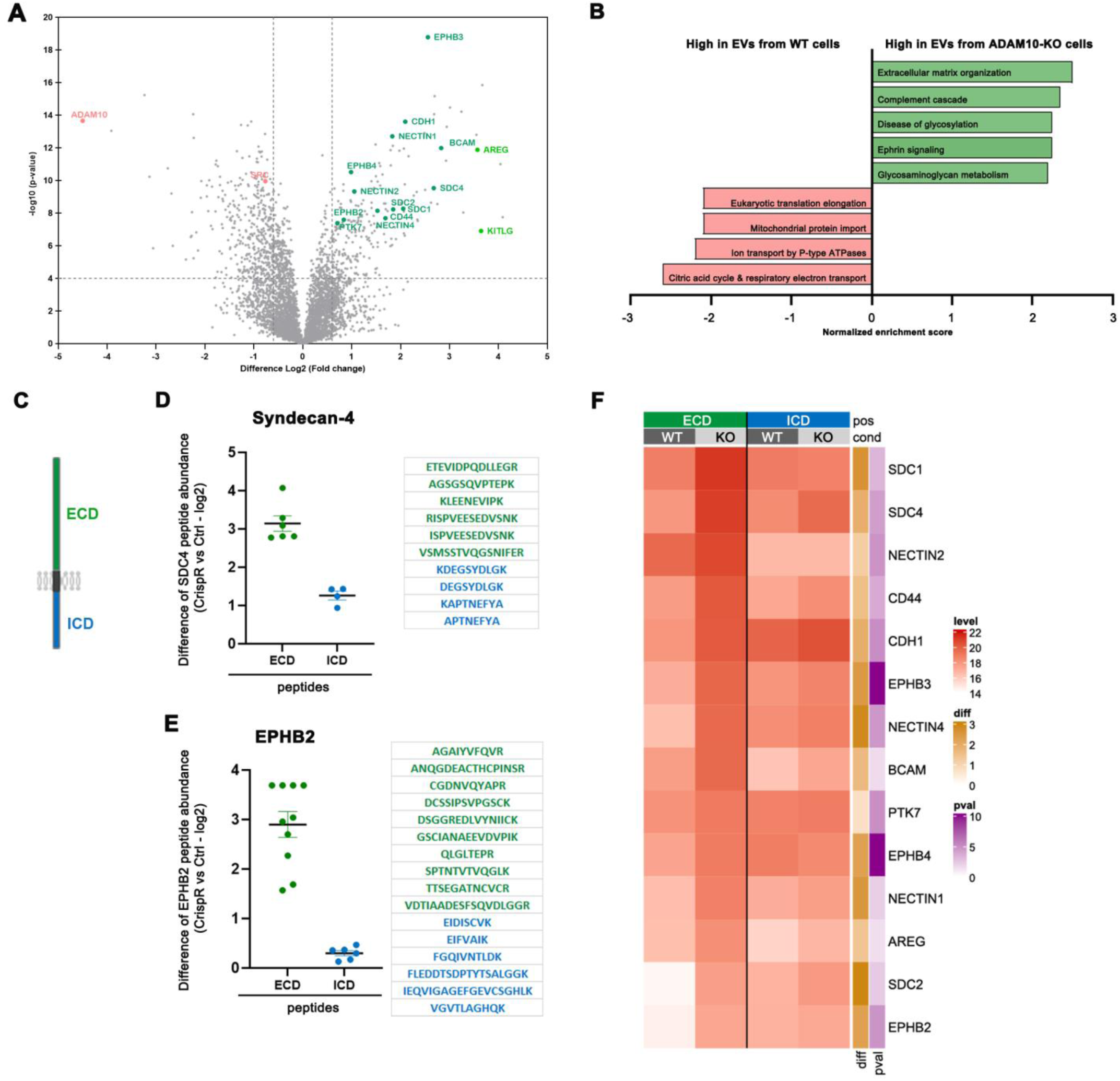
ADAM10 knock-out impacts sEV protein composition and increases the loading of receptor ectodomains / non-cleaved receptors into sEVs. sEVs secreted by MCF7 cells with knock-out of ADAM10 (ADAM10-KO) or control cells (WT) were isolated by differential ultracentrifugation from conditioned media and analyzed by mass spectrometry. **A**. The volcano plot illustrates the proteins that are differentially expressed, with log2 levels (x-axis) and −log10 (P-value) (y-axis), in ADAM10-KO versus WT sEVs. Proteins downregulated (left) or upregulated (right) in ADAM10-KO were selected based on a p-value < 0.0001 and a difference > 1.5. **B.** Functional annotation was performed through Gene Set Enrichment Analysis (GSEA) using Webgestalt based on pathway/reactome data. Note that extracellular matrix proteins are enriched in sEV ADAM10-KO (green) while intracellular proteins (red) are enriched in sEV from WT cells. **C.** Schematic representation of a transmembrane type I protein illustrating the color code used under D-E for peptides as identified by mass spectrometry. **D-E** The difference (y-axis) of log2 peptides abundance obtained by mass spectrometry analysis in ADAM10-KO sEVs compared to control WT sEVs is represented for SDC4 (SDC4, **D**) and Ephrin-B2 (EPHB2, **E**). The extracellular domain (ECD) peptides are represented in green, and the intracellular domain (ICD) peptides are in blue. **F.** Heat map illustrating the level (log2 of abundance, gradient from white to red) of ECD (green) or ICD (blue) peptides in sEVs from WT or ADAM10-KO cells for different receptors (indicated on the right). The difference (diff, shown in ochre) corresponds to the difference between ECD and ICD peptides of the same receptor, based on their respective abundance difference between KO and Ctrl conditions (y-axis in panels D and E). The P-value of this difference (obtained by an unpaired two-tailed t-test) is presented as −log10(P-value) (pval in magenta). pos = peptide position on the receptor, cond = condition.

### ADAM10 downregulation supports the incorporation of full-length membrane-associated signaling receptors into sEVs

While proteomic data confirmed that SDC ectodomains/full-lengths forms are more abundant in ADAM10-KO sEVs (see Fig. 3C-D for SDC4), it was also striking, that the same holds for other signaling receptors. For example, the peptides corresponding to the extracellular domain of EPHB2 were significantly enriched in ADAM10-KO sEVs (Fig. 3E). In fact, upon ADAM10 loss sEVs were enriched in peptides corresponding to the extracellular domain of a plethora of signaling receptors (Fig. 3F). Western-blot analyses confirmed that the full-length forms of EphrinB3, BCAM, E-cadherin and PTK7 are enriched in ADAM10-KO sEVs (Fig. S4B), while the intracellular tyrosine kinase SRC is downregulated (Fig. S4C). Importantly, the cell lysate levels of these proteins were unchanged (Fig. S4B). Together, these data demonstrate that ADAM10 shapes not solely sEV composition, but also EV surface or corona.

### Impact of ADAM10 on sEV cytosolic delivery

In the light of the message conveyed by the proteomic data, we further investigated whether ADAM10 may impact sEV mode of signaling - either internal cargo delivery and/or contact signaling.

To evaluate whether ADAM10 may impact the ability of sEVs to deliver their internal content into the cytoplasm of recipient cells, we used a split-Nanoluciferase (NL) assay based on the system described by Dixon et al. (Dixon *et al*, 2016) and as further developed by Hyka et al. (Hyka *et al*, 2026). This setup involves preparation of sEVs by concentration of conditioned medium from HEK293 cells containing HiBiT-syntenin and MCF7 recipient cells expressing LgBiT in their cytosol (Fig. 4A). Here, e compared the ability of HiBit-syntenin sEVs originating from MCF7 cells, inhibited or not for ADAM10, to reconstitute Nanoluciferase in recipient MCF7 LgBiT cells. First, we observed the capture of HiBiT-syntenin sEVs by cells was comparable (Fig. 4B). Yet, while sEVs from ADAM10-high cells were prone to deliver HiBiT-syntenin in the cytoplasm of MCF7 recipient cells, sEVs from ADAM10-inhibited MCF7 cells clearly failed to deliver their cargo, with signals below background levels (Fig. 4C). Similar results were obtained with HiBiT-syntenin sEVs originating from ADAM10-inhibited HEK293 cells (Fig. S5C). Of note, ADAM10 inhibition affected sEV composition in HEK293 cells similarly as in other cells (Fig. S5A-B). These data indicate that ADAM10 activity in sEV donor cells may facilitate effective cytosolic delivery of sEV content.

**Figure 4.**
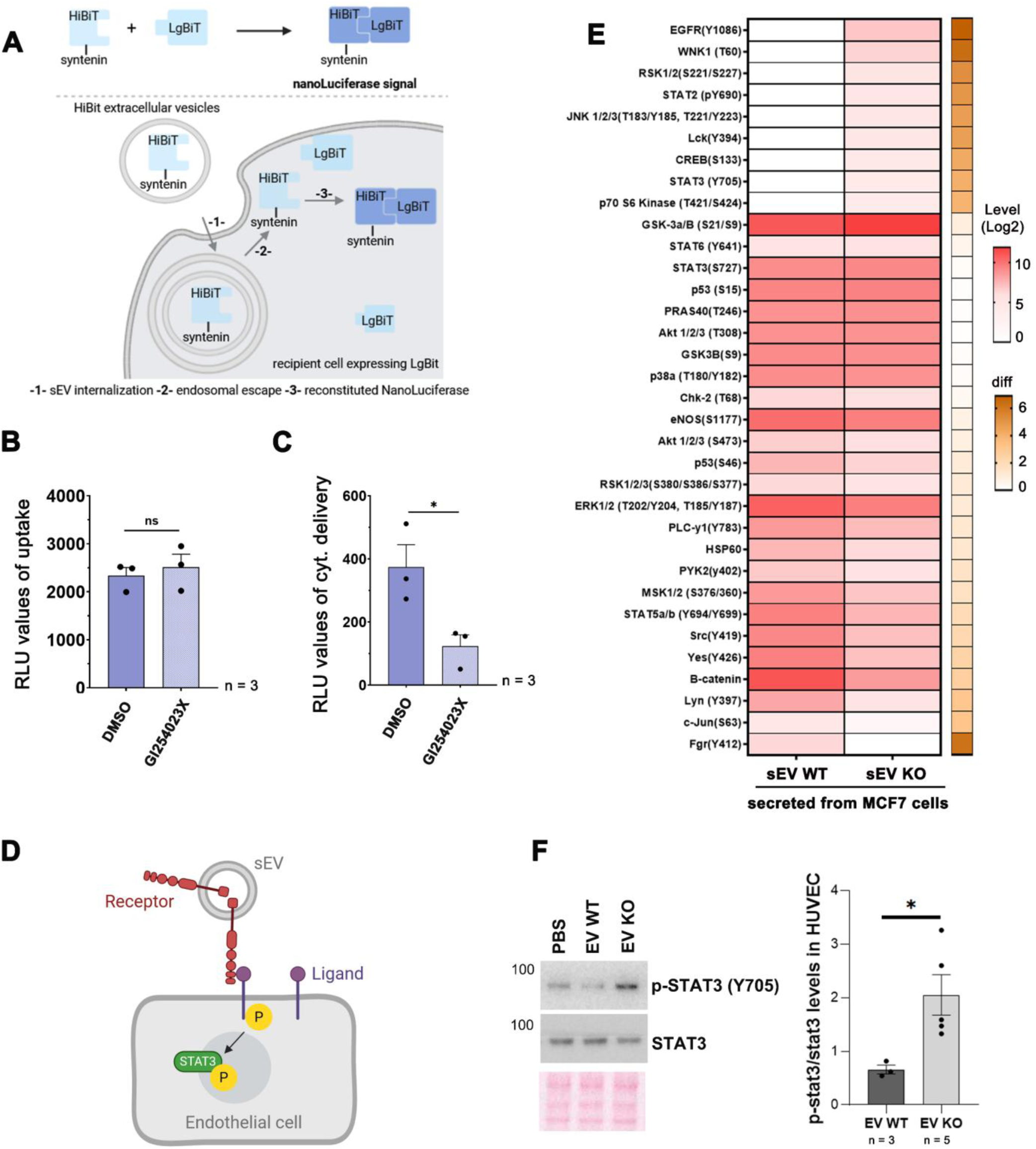
ADAM10 modulates the sEV signaling. **A.** Principle of the endosomal escape assay. Nanoluciferase can be split into two non-functional fragments (HiBiT and LgBiT) that reconstitute enzymatic activity upon interaction. In this setup, HiBiT is fused to syntenin and thus is loaded into sEVs from HEK293 cells. HiBiT-syntenin sEVs (concentrated conditioned medium downregulated for the large EV fraction by centrifugation at 10.000 g) are incubated for 4hours with MCF7-recipient cells that overexpress LgBiT. After HiBiT-syntenin sEVs are taken up (1), nanoluciferase will be reconstituted solely if HiBiT-syntenin sEVs undergo endosomal escape (2), and liberate HiBiT-syntenin in the cytosol to associate with LgBiT (3). **B.** Histogram illustrating that the uptake of HiBiT-syntenin sEVs by MCF7 LgBiT cells after 4 hours of incubation (1 in A) is comparable regardless of treatment with the ADAM10 inhibitor GI254023X. **C.** Histogram illustrating that endosomal escape of HiBiT-syntenin sEVs (2 in A) is abolished when sEVs originate from MCF7 treated with ADAM10 inhibitor (GI254023X). Histograms represent mean signal intensities in relative light units (RLU) after addition of the nanoluciferase substrate Furimazine, ± SEM. * P < 0.05, n.s. non-significant (one-way ANOVA). n indicates the number of independent experiments; each point represents one independent experiment. Dashed line corresponds to background signals. **D.** Scheme illustrating how non-cleaved receptors present on sEVs can bind corresponding ligands on endothelial cells and induce STAT3 phosphorylation. **E.** Heatmap showing log 2-fold changes of protein phosphorylation after treatment with sEV from MCF7 ADAM10 KO cells compared to control WT cells. The level of average densitometric values (Log2) are presented as a spectrum of color where white and red colors represent the lowest and the highest values in the row according to the indicated scale. The difference (diff, shown in ochre) corresponds to the difference between KO and WT conditions of the same protein, based on their respective signal intensity difference between KO and Ctrl conditions. The experiment was performed once. **F.** Representative Western blots illustrating the levels of phosphorylated STAT3 (p-STAT3) and total STAT3 in cell lysates from HUVECs treated for 30 minutes with PBS, sEV secreted by MCF7 WT cells (sEV WT) or sEV secreted by MCF7 ADAM10-KO (sEV KO) cells, as indicated. sEVs were prepared by size exclusion chromatography. Histogram represents mean signal intensities of p-STAT3/STAT3 normalized to signals in control (PBS) ± SEM calculated from 3 independent experiments. * P < 0.05 (unpaired non-parametric Mann–Whitney test).

### Impact of ADAM10 on sEV signaling by contact

Since sEVs from ADAM10-KO cells are enriched in several full-length membrane proteins (Fig. 3C-F), we hypothesized that sEVs from ADAM10-downregulated donor cells may more effectively engage their cognate ligands on sEV recipient cells, thereby promoting receptor activation and subsequent downstream recruitment and phosphorylation of key transcription factors (Fig. 4D). To use an *in vitro* model relevant in the tumoral context, we used HUVEC endothelial cells as recipient for our breast cancer cell sEVs. We further identified proteins that become phosphorylated using a phospho-array that can detect up to 37 different phosphorylated proteins. sEVs secreted by WT and ADAM10-KO MCF7 cells were isolated using size exclusion chromatography (Fig. S5D left), to preserve sEV fine structure. Changes in phosphorylated signal intensities in HUVEC cells following treatment with WT or ADAM10-KO sEVs are presented as heat maps (Fig. 4E and Fig. S5E). Phosphorylated proteins in HUVEC cells were also investigated after treatment with sEVs secreted by WT and ADAM10-KO MDA-MB-468 cells (Fig. S5D, right and S5F). Among the monitored proteins, 9 and 13 proteins showed increased phosphorylation in response to ADAM10-KO sEVs from MCF7 and MDA-MB-468, respectively with 5 proteins (EGFR, WNK1, STAT2, CREB and STAT3) shared between both conditions. Western blot analysis further validated a two-fold increase in STAT3 Y705 phosphorylation in HUVEC cells treated with ADAM10-KO sEVs compared to PBS or WT sEVs. (Fig. 4F). Altogether these results indicate that when ADAM10 is inhibited in breast cancer sEV donor cells, those EVs are more prone to activate signaling pathways known to be contact-dependent.

## Discussion

We here identified solely two proteases associated with SDC immunoprecipitates. This was unexpected as many other proteases are expressed by MCF7 cells. Surprisingly, although ADAM10 and ADAM17 are typically considered to cleave overlapping substrates (Pruessmeyer & Ludwig, 2009), our data demonstrate that solely ADAM10 affects SDC processing in MCF7 cells and their sEVs. Given that previous studies reported ADAM17-mediated cleavage of SDCs (Pasqualon *et al*, 2015; Pruessmeyer *et al*, 2010), and that both ADAM10 and ADAM17 associated to EVs can contribute to substrate shedding (Groth *et al*, 2016; Stoeck *et al*, 2006), the lack of involvement of ADAM17 in this study may reflect cell-type-specific regulation. Such may be related to differential expression of proteins like tetraspanins (Yáñez-Mó *et al*, 2011), which are known to influence protease targeting and activity (Jouannet *et al*, 2016; Saint-Pol *et al*, 2017). Noteworthy, a similar role for ADAM10 was consistently observed across all four cell types tested in this study, indicating that regulation of sEV-SDC by ADAM10 may be quite generic. We also observed that ADAM10 did not influence the number of secreted EVs in any of these cell types tested. So, our data are not directly supportive of a recent study (Bizingre *et al*, 2025) suggesting that ADAM10 is involved in the detachment of sEV from the cell surface in neuroepithelial cells.

Our findings indicate a critical role of SDCs in controlling the incorporation of ADAM10 in sEVs. In the absence of SDCs, ADAM10 accumulates in cell lysates and in proximity with CD63-positive endosomal compartments. As SDC and ADAM10 co-immunoprecipitate, a plausible scenario is that SDC hooks ADAM10 to syntenin and ALIX for intraluminal budding into multivesicular bodies. Yet, it is striking that ADAM10, regulating SDC processing, and drastically influencing sEV composition does not alter syntenin, ALIX or tetraspanins abundance in sEVs. Of note, although ADAM10 knock-down markedly influences the ratio of full-length (FL) versus C-terminal fragment (CTF) forms of SDC in cells and sEVs, it does not influence the net amount of SDC-ICD in sEVs. These observations lend support to the concept that SDC-syntenin-ALIX function as a core machinery for the biogenesis of sEVs (Baietti *et al*, 2012; Castro-Cruz *et al*, 2023). More importantly, this study offers an important new level of understanding of the core mechanisms controlling sEV biology (Figure 5). In particular, the accumulation of glycosaminoglycans (GAG)-substituted FL SDCs in sEVs from ADAM10-KO cells support the notion that ADAM10 downregulation may redirect sEV budding towards the plasma membrane (Figure 5, upper left). This because the large size and the strong repulsive negative charge imposed by these GAG-structures renders the double-membrane budding process characteristic of sEV biogenesis within MVBs biophysically improbable. When ADAM10 is active, SDC-syntenin-ALIX can bud within MVBs for the production of sEVs, and SDC is cleaved by ADAM10 during the process, explaining why it accumulates in sEV as CTF (Figure 5, lower left). Cleavage by ADAM10 may require prior GAG-removal by heparanase (Roucourt *et al*, 2015; Lambaerts *et al*, 2009). Such scenario is also totally in line with the change of sEVs composition contain more cytosolic proteins when ADAM10 is active and may also imply that proteins may not be equally distributed from the cell interior to the cell surface (Figure 5, see red sEVs and green sEVs alluding to their different composition according to their budding site).

**Figure 5.**
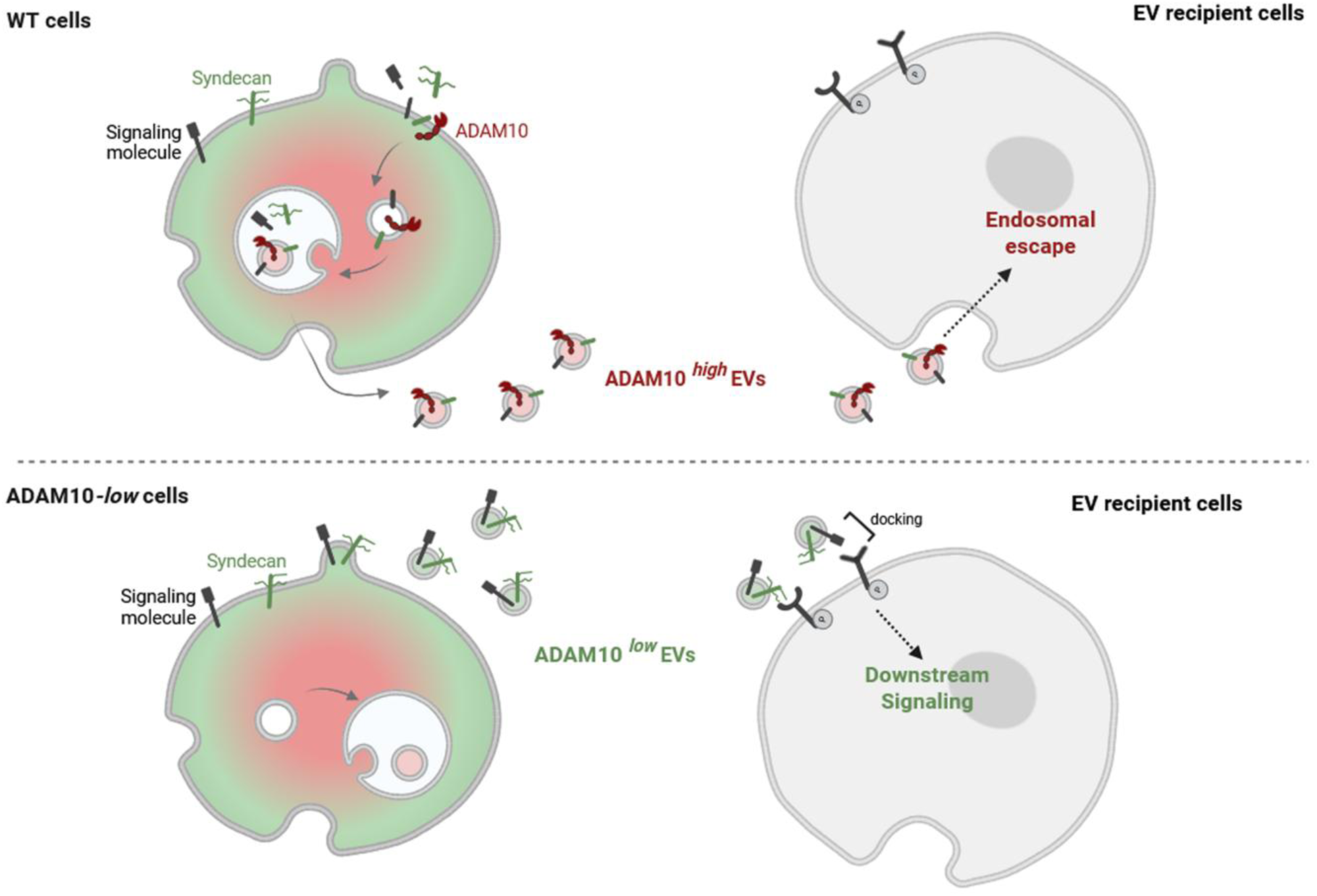
Current model to recapitulate the findings of the present study. sEVs originating from ADAM10*^low^* cells (lower panel) are decorated with non-cleaved receptors (including SDC-FL) and support signaling by contact as illustrating for STAT3 phosphorylation in HUVEC cells. sEVs originating from ADAM10*^high^* cells (upper panel) are decorated with cleaved receptors (including SDC-CTF) and support the endosomal escape of internal cargo’s as shown for syntenin. sEVs originating from ADAM10*^high^* cells are also more enriched in cytosolic component while sEVs originating from ADAM10*^low^* cells are more enriched in membrane signaling component. As cellular content is not highly different between ADAM10*^high^* and ADAM10*^low^* cells, and as the presence of GAG-substituted SDC on sEVs is not compatible with the double membrane budding process characteristic of sEV originating from MVB, we propose that sEV bud from the plasma membrane when ADAM10 is inactive and from MVB when ADAM10 is active. See text for more details. This figure was created using BioRender.com

Although in the context of SDC-dependent sEV production, we favor a scenario where ADAM10 proteolytic cleavage occurs intracellularly at the level of endosomes, we did not directly determine whether ADAM10 cleaves its many substrates intracellularly or on sEVs after release. In fact, both mechanisms are plausible and could co-exist. Notably, it has been shown that ADAM10-mediated cleavage of CD23, the receptor for immunoglobulin E, occurs after its incorporation into EVs but prior to their release from the cell (Mathews *et al*, 2010). Moreover, even though EGFR ectodomain shedding can occur both in cells or in EVs after their release, EGFR shedding occurs predominantly inside the cell (Sanderson *et al*, 2008). Of note, the presence and enzymatic activity of ADAM10 in circulating EV have been extensively documented (Sharples *et al*, 2008; Tosetti *et al*, 2018; Lin *et al*, 2019; Rai *et al*, 2025) in pathological contexts. For example in cancer, ADAM10 in EVs was identified as a potential breast cancer biomarker (Risha *et al*, 2020). In addition, in the context of Alzheimer’s disease, it has been proposed that APP cleavage may occur within EVs enriched in proteases such as ADAM10 (Sharples *et al*, 2008). Importantly, ADAM10 activity within EVs can mediate the release of cytokines and growth factors that may interfere with immunotherapy (Tosetti *et al*, 2018). Finally, cleavage could even occur *in trans* as human cancer-associated fibroblasts were shown to secrete ADAM10-enriched sEVs that activate Notch signaling in recipient cancer cells (Shimoda *et al*, 2014).

Another key finding of this study is that ADAM10 supports the formation of sEVs competent to deliver their internal content after internalization in recipient cells – documented for syntenin (Figure 5, bottom right). The absence of ADAM10, on the contrary compromise such delivery and supports the formation of sEVs containing non-cleaved receptors on their surface. In contrast to cleaved forms, these full-length receptors enable direct contact-dependent signaling with recipient cells. In particular, ADAM10 sEVs enriched in specific non-cleaved receptors significantly modulate downstream signaling in endothelial target cells. It would be interesting to better understand whether surface/adhesion molecules co-localize on the same sEV subtypes or are distributed across distinct syndecan-FL-derived sEV populations. This will refine our understanding of the functional heterogeneity within the sEV pool and may uncover specific vesicle subtypes that mediate unique biological effects.

In conclusion, our results identify ADAM10 as a key regulator of sEV identity and function, shaping EV surface architecture and internal cargo delivery, and thereby influencing their signaling modes and biological activity. These findings provide a framework for future studies exploring how the modulation of surface molecule processing in sEVs can be exploited for diagnostic or therapeutic purposes. Furthermore, we demonstrate that ADAM10 incorporation into sEVs is SDC-dependent, highlighting SDCs as potential targets to fine-tune receptor cleavage in various physio-pathological contexts.

---

**Figure S1.**
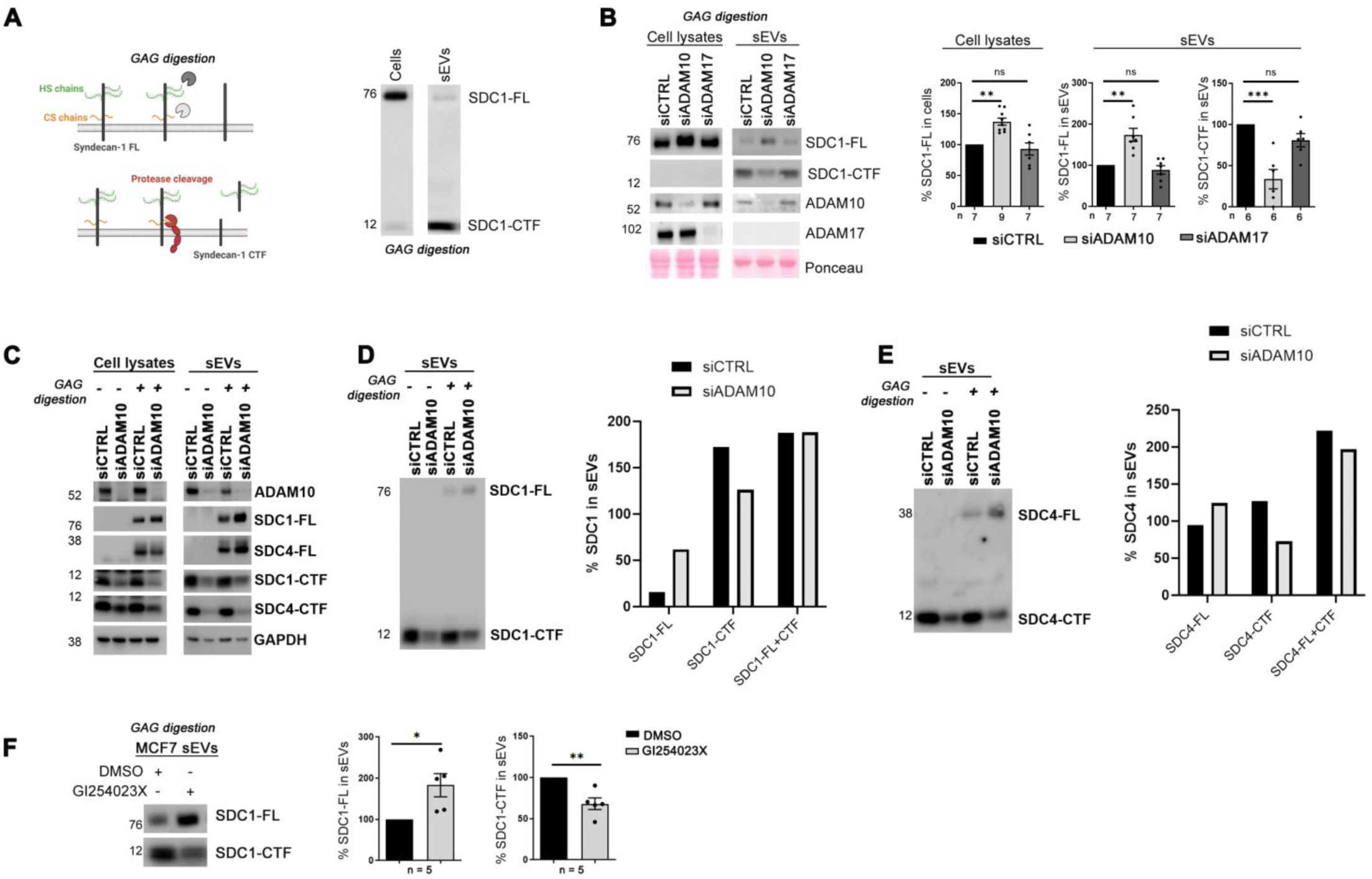
ADAM10 (but not ADAM17) downregulation or inhibition modifies sEV-SDC1. **A**. **Left:** Scheme illustrating the complexity of SDC1 structure and processing. (Upper part) SDC1 full-length (FL) core protein (black) is substituted with glycosaminoglycan chains (GAG) of the heparan (green) and chondroitin (orange) sulfate type. The trimming of GAG chains by specific enzymes is necessary for the FL protein to migrate at a discrete band in SDS-PAGE and to be detectable by immunoblotting. (Lower part) SDC1 core protein can be cleaved by proteases generating two main fragments: an N-terminal fragment comprising most of the GAG-substituted extracellular domain (ECD) and a C-terminal fragment (CTF) comprising the remainder of the ECD, the membrane-spanning and the cytoplasmic domain. **Right:** Western blot illustrating the signals obtained, after GAG-digestion, for FL SDC1 and SDC1 CTF in the cells and the sEV enriched fraction obtained after differential ultracentrifugation of the conditioned extracellular media. Signals were obtained with an antibody recognizing the intracellular domain of SDC1. Note that the FL form of SDC1 (SDC1-FL) abounds in cell lysates, while the CTF is less abundant. On the contrary, the SDC1-CTF is abundant and the SDC1-FL is barely detectable in sEVs. **B.** MCF7 cells downregulated for ADAM10 (siADAM10) or ADAM17 (siADAM17) and control cells (siCTRL) were evaluated for SDC1 FL and CTF abundance in cells and sEVs by Western blot, after GAG-digestion. Histograms represent the mean signal intensity for indicated proteins relative to the signal in control cells, ± SEM. Statistical analysis was performed using the Kruskal-Wallis one-way non-parametric ANOVA test (* P < 0.05, ** P < 0.01, *** P < 0.001, **** P < 0.0001, n.s. non-significant). **C.** MCF7 cells downregulated for ADAM10 (siADAM10) and control cells (siCTRL) were treated with heparitinase and chondroitinase (GAG digestion +) or not (GAG digestion -) to evaluate SDC substitution with GAG chains. SDCs FL and CTF in cells and sEVs were analyzed by Western blot as indicated. Single blots examining the relative abundance of FL versus CTF forms of SDC1 (**D**) or SDC4 (**E**) forms are provided. **F**. sEVs secreted by MCF7 cells inhibited for ADAM10 activity, (GI254023X) versus controls (DMSO) were isolated from conditioned media by differential ultracentrifugation. sEVs were analyzed by Western blot after GAG-digestion to evaluate the levels of SDC1 FL and CTF. Histograms represent the mean signal intensity for indicated proteins relative to the signal in control cells, ± SEM. * P < 0.05, ** P < 0.01, *** P < 0.001, **** P < 0.0001, n.s. non-significant (unpaired non-parametric Mann–Whitney test). n indicates the number of independent experiments; each point represents one independent experiment.

**Figure S2.**
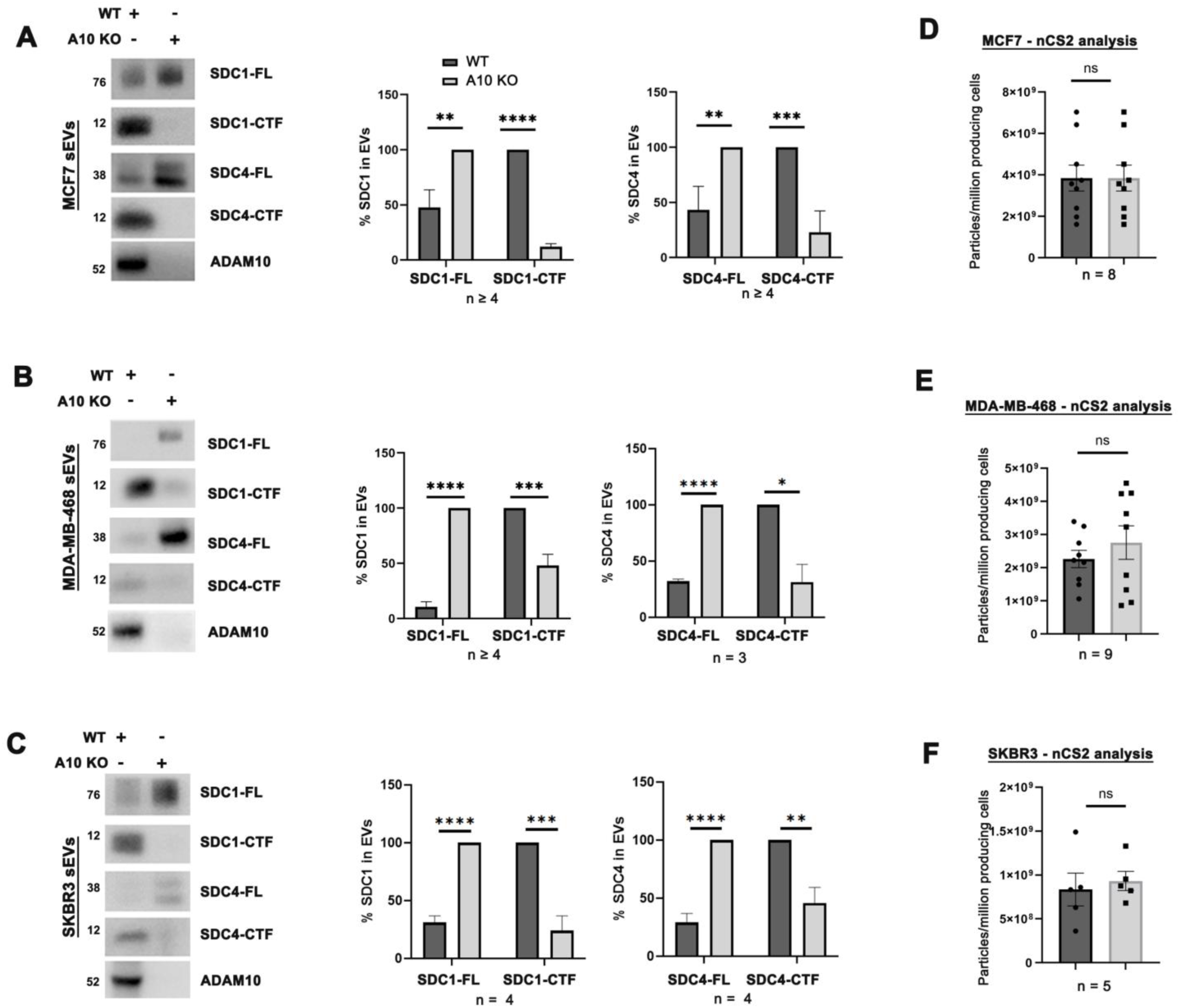
ADAM10 knock-out modifies sEV-associated -SDC1 and -SDC4 in various cell types. sEVs secreted by MCF7 (**A, D**) MDA-MB-468 (**B, E**) or SKBR3 (**C-F**) cells with knock-out of ADAM10 (A10 KO) or control cells (WT) were isolated from conditioned culture media by differential ultracentrifugation. sEVs were analyzed by Western blot after GAG-digestion to evaluate the levels of SDC1 and SDC4 FL and CTF. Histograms under **A-C** represent the mean signal intensity for indicated proteins relative to the signal in ADAM10-KO cells for SDC-FL and to the signal in control cells for SDC-CTF, ± SEM. The value ‘n’ indicates the number of independent experiments. * P < 0.05, ** P < 0.01, *** P < 0.001, **** P < 0.0001, n.s. non-significant (multiple unpaired t-test). **D-F.** The number sEVs secreted by MCF7 (**D**), MDA-MB-468 (**E**), SKBR3 (**F**) cells knock-out for ADAM10 (ADAM10-KO) or control cells (WT) were further analyzed by microfluidic resistive pulse sensing (MRPS) - Spectradyne (nCS2). n indicates the number of independent experiments.

**Figure S3.**
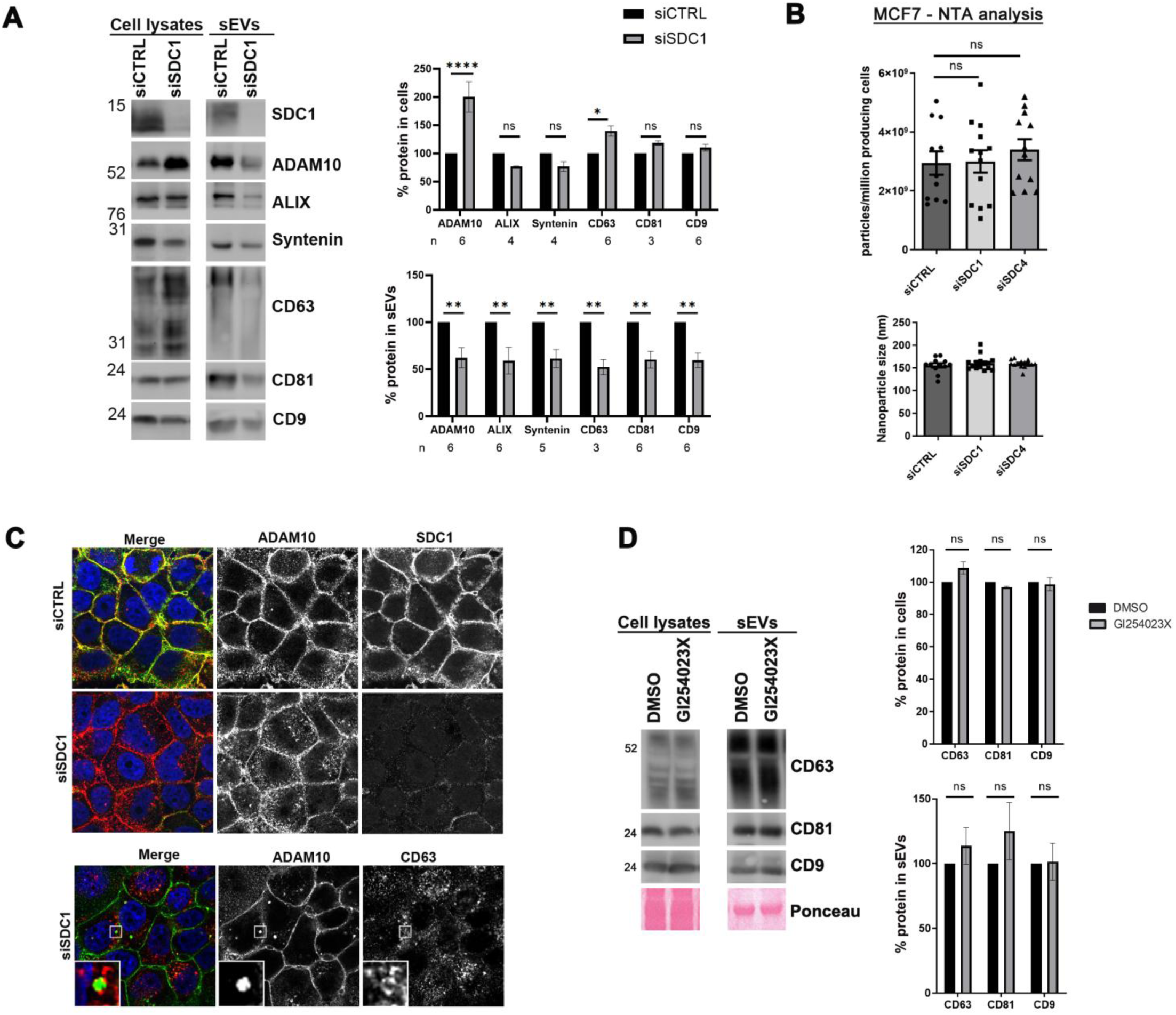
ADAM10 loading in sEVs is SDC1-dependant and ADAM10 downregulation does not change syntenin, ALIX and tetraspanins (CD9, CD63, CD81) in sEVs. **A.** sEVs secreted by SDC1-downregulated cells (siSDC1) versus controls (siCTRL) were isolated by differential ultracentrifugation. sEVs were further analyzed by Western blot, testing for several markers, as indicated. Histograms represent mean signal intensities in sEVs relative to signals obtained for controls, ± SEM. * P < 0.05, ** P < 0.01, *** P < 0.005, **** P < 0.001, n.s. non-significant (two-way ANOVA). **B.** sEVs secreted by MCF7 cells downregulated for SDC1 (siSDC1), SDC4 (siSDC4) or control cells (siCTRL) were isolated by differential ultracentrifugation from conditioned media and further analyzed by Nanosight (NTA) to determine concentration and size, as indicated. Each point represents one independent experiment. **C.** (Upper part) Representative confocal micrographs showing the steady-state distribution of endogenous ADAM10 (red in merge) in MCF7 cells transfected with control (siCTRL) siRNA or siRNA targeting SDC1 (siSDC1). SDC1 was stained with antibody recognizing intracellular domain of SDC1 (green in merge). (Lower part) Representative confocal micrographs showing the steady-state distribution of endogenous ADAM10 (green in merge) together with CD63 (red in merge) upon SDC1 downregulation. In merge, nuclei are stained with DAPI (blue). **D.** sEVs secreted by cells treated with the ADAM10 inhibitor GI254023X versus DMSO controls were isolated by differential ultracentrifugation. Cell lysates from parent cells or sEVs were further analyzed by Western blot, testing for several markers, as indicated. Note that ADAM10 inhibition does not impact the loading of tetraspanins CD63, CD81 and CD9. Histograms represent mean signal intensities relative to signals obtained for controls, ± SEM. * P < 0.05, ** P < 0.01, *** P < 0.001, n.s. non-significant (two-way ANOVA). The value ‘n’ indicates the number of independent experiments.

**Figure S4.**
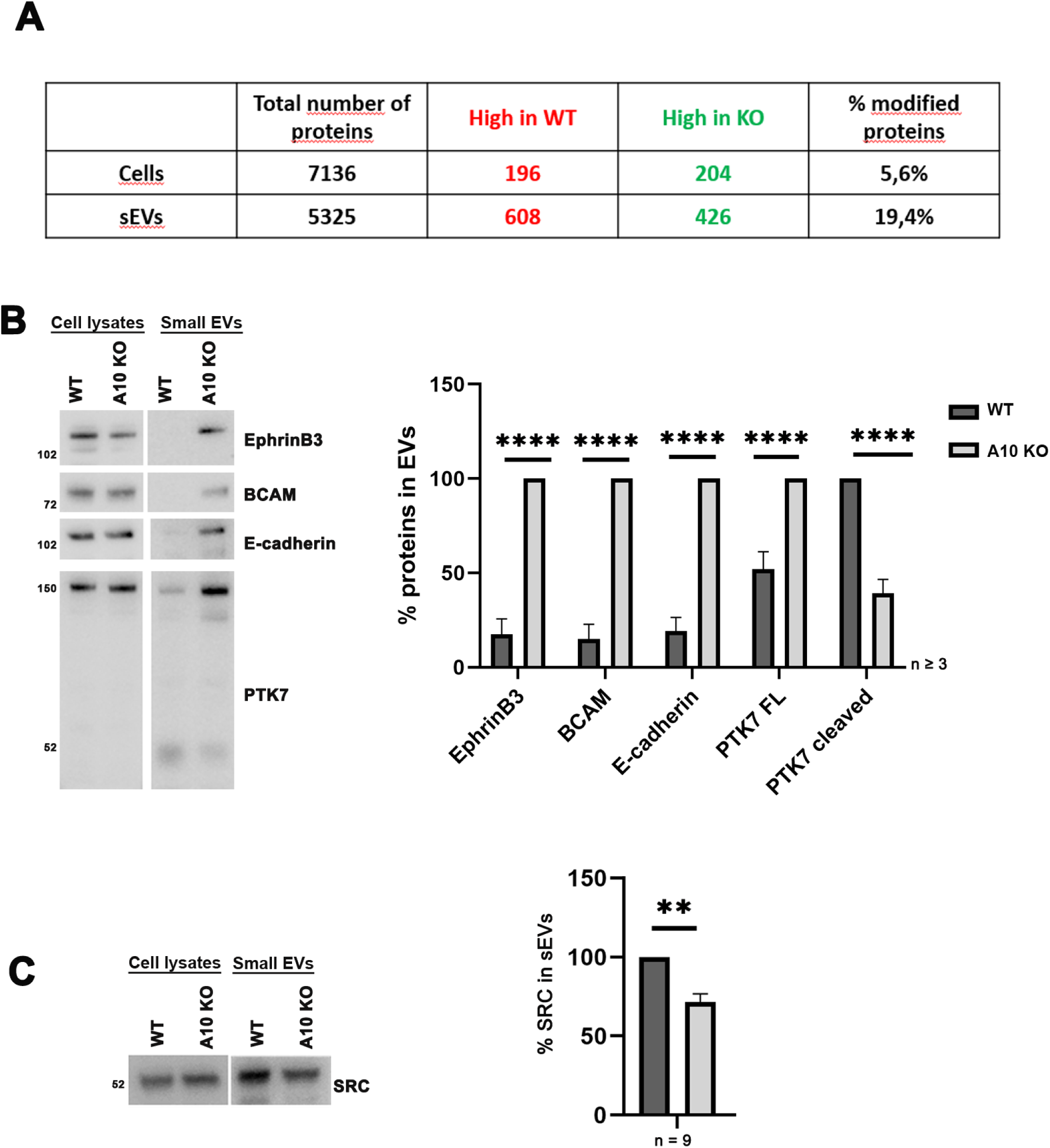
Validation in Western blot of the changes of some sEV cargoes in WT and ADAM10-KO conditions. **A.** Summary table presenting the proteins identified and quantified by mass spectrometry analysis of ADAM10-KO cell lysates compared to control MCF7 cells and their corresponding secreted sEVs. **B-C.** sEVs secreted by MCF7 cells with knock-out of ADAM10 (A10 KO) or control cells (WT) were isolated by differential ultracentrifugation of conditioned culture media. Cell lysates and sEVs were analyzed by Western blot, testing for several proteins including EphrinB3, BCAM, E-cadherin and PTK7 (**B**), SRC (**C**). Histograms represent mean signal intensities normalized to controls ± SEM, calculated from n independent experiments. * P < 0.05, ** P < 0.01, *** P < 0.001, **** P < 0.0001, n.s. non-significant (multiple unpaired t-test).

**Figure S5.**
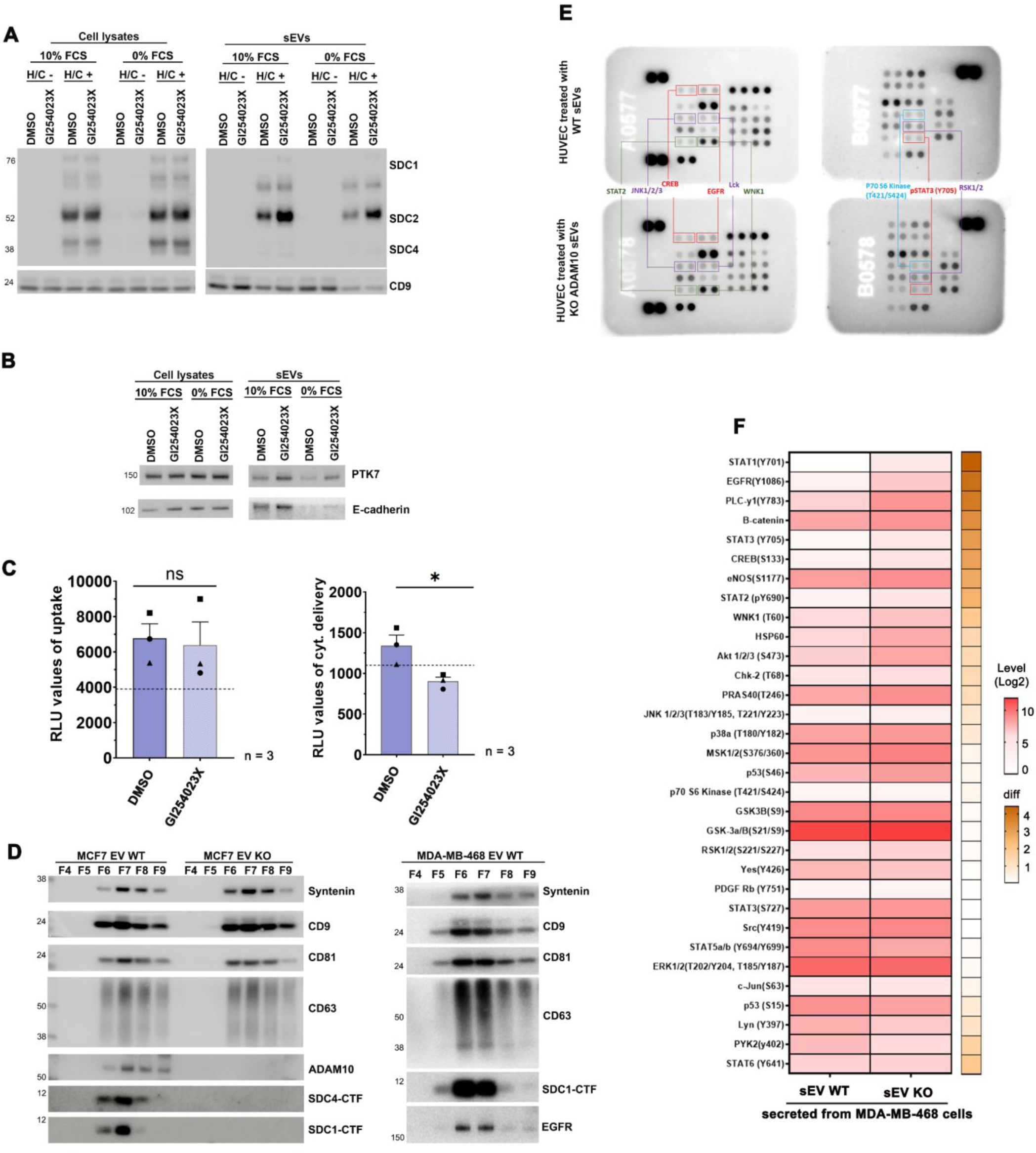
**A-B.** Western blot of the cell lysates and sEVs from HEK293 cells treated with the ADAM10 inhibitor GI254023X or DMSO as control, in the presence or absence of serum (FCS), show that ADAM10 supports the cleavage of various receptors, as indicated by the observed increase of non-cleaved receptors in the sEV of those cells, irrespectively of serum (although the absence of serum significantly reduces sEV accumulations). These control experiments were performed to ensure the pertinence of the endosomal escape assay as developed by Hyka et al., 2025 in the context of the present study. For SDCs (A), cells were treated with heparatinase and chondroitinase (GAG digestion +) or not (GAG digestion -). Heparan sulfate proteoglycans / SDCs are detected with the mAb 3G10 recognizing GAG chain stubs left after digestion (David *et al*, 1992). **C.** Left. Histogram illustrating that the uptake of HiBiT-syntenin sEVs by HEK293 LgBiT cells after 4 hours of incubation is comparable regardless of treatment with the ADAM10 inhibitor GI254023X. Right. Histogram illustrating that endosomal escape of HiBit-syntenin sEVs is abolished when sEVs originate from HEK293 treated with ADAM10 inhibitor (GI254023X). Histograms represent mean signal intensities in relative light units (RLU) after addition of the nanoluciferase substrate Furimazine. **D.** Illustrative western blot of size exclusion chromatography (SEC) experiments for the preparation of sEVs for functional assays. Left. Syntenin, CD9, CD81, CD63, ADAM10, SDC1 and SDC4 were used as markers to select ad-hoc fractions from the conditioned media of MCF7 cells KO for ADAM10 (sEV KO) or control cells (sEV WT) as indicated. Fractions 6-9 were pooled before incubation with HUVEC cells. Right. Syntenin, CD9, CD81, CD63, SDC1 and EGFR were used as markers to select ad-hoc fractions from the conditioned media of MDA-MB-468 WT cells as indicated. Fractions 6-8 were pooled before incubation with HUVEC cells. **E.** The phosphorylation of selected proteins (top nine proteins showing increased phosphorylation after ADAM10-KO EV treatment) were determined using the phospho-array (R&D systems). Signals for each phosphorylated protein are presented as a pair of duplicate spots for the same exposition of 15 minutes. Average densitometric values for the phosphorylated proteins are shown in the heatmap Fig. 4E. **F.** Heatmap showing log 2-fold changes of protein phosphorylation after treatment with sEV from MDA-MB-468 ADAM10 KO cells compared to control WT cells. The log 2-fold changes of protein phosphorylation after sEV KO treatment compared to control (sEV WT) were shown in the heatmap. The level of average densitometric values (Log2) are presented as a spectrum of color where white and red colors represent the lowest and the highest values in the row according to the indicated scale. The difference (diff, shown in ochre) corresponds to the difference between KO and WT conditions of the same protein, based on their respective signal intensity difference between KO and Ctrl conditions. The experiment was performed two or three times depending on the analyzed protein.

## Materials and Methods

### Cells and transfections

MCF-7, MDA-MB-468, SKBR3 and HEK293 cells were obtained from ATCC (Manassas, VA). Cells were routinely grown in DMEM medium (Thermo Fisher Scientific,) supplemented with 10% fetal calf serum (FCS) (Sigma Aldrich). HUVEC cells were a kind gift from Michel Aurrand-Lions, CRCM Inserm UMR1068, Marseille, France. HUVEC cells were routinely grown in Endothelial Cell Growth Medium 2 (Promocell) supplemented with Endothelial Cell Growth Medium 2 supplement mix (Promocell). Serum used for experiments was downregulated of EVs, by prior overnight centrifugation at 110,000 x g. For transient expressions, the cells were transfected the day after plating using Fugene HD reagent (Roche). For RNAi experiments, cells at a confluence of 50% were transfected with 20nM RNAi using Lipofectamine RNAiMAX reagent (Life technologies, USA); cells were analyzed 48 or 72 hours after RNAi treatment. MCF7, MDA-MB-468 and SKBR3 cells KO for ADAM10 were generated using CRISPR/Cas9 technology. The sgRNA oligos targeting ADAM10 were cloned into the lentiCRISPRv2 plasmid (generous gift of E. Rubinstein).

### Expression vectors and reagents

For gain-of-function studies in cells, the cDNA encoding full-length GFP-SDC4 was provided by C. Addi (Addi *et al*, 2020). All RNAi targeting human sequences (siGENOME) and the non-targeting control RNAi (siCTRL) were purchased from Horizon Discovery. ADAM10 (5’ GGACAAACTTAACAACAAT 3’), ADAM10 SMARTpool (M-004503-02), ADAM17 SMART pool (M-003453-01), SDC1 (5’ CGCAAAUUGUGGCUACUAA 3’), SDC4 (5’ GUGAGGAUGUGUCCAACAA 3’). The GI254023X inhibitor (used at 5 µM) was purchased from Sigma Aldrich (SML0789).

### Western blotting

Cells were lysed in 30 mM Tris-HCl (pH 7.4), 1% NP40, 150 mM NaCl, 1 mM EDTA, 1 _μ_g/ mL Pepstatin, 1 µg/mL Leupeptin (Sigma-Aldrich). sEVs were suspended in PBS. Four volumes of lysates/suspension were mixed with one volume 5x Laemmli sample buffer (250 mM Tris-HCl pH 6.8, 25% glycerol, 10% SDS, bromophenol blue). The heat-denaturated proteins were fractionated in 4-12 % gels by SDS–PAGE, electro-transferred to nitrocellulose membrane, stained with Ponceau red, and immunoblotted with the indicated antibodies. Signals were visualized using chemiluminescence HRP substrate ECL kit (Amersham ECL Detection Reagents) and Amersham ImageQuant 800 imaging system (Cytiva). Quantification was performed by densitometry using ImageJ.

Anti-syntenin and anti-ALIX antibodies were as described earlier (Baietti *et al*, 2012). ADAM10, CD63, CD81 and CD9 antibodies were provided by E. Rubinstein (Sorbonne University, CIMI-Paris). Two peptides corresponding to the C-terminal fragment of SDC4 (DLGKKPIYKKAPTNE and DLGKKPIYKKAPTNEFYA) were used to generate SDC4-rabbit polyclonal antisera (Eurogentec). Other antibodies were from commercial sources and were used as recommended by the manufacturers. Antibodies directed against SDC1 intracellular domain (D4Y7H) was from (cell signaling #12922, dilution 1/1000), GFP (A11122) from Thermofisher (dilution 1/1000), ADAM17 (abcam, #ab39162, 1/1000 or cell signaling #3976, 1/1000), ADAM10 (abcam, #ab1997, 1/1 000), GAPDH (Proteintech # 10494-1-AP), Flotillin-1 (BD Biosciences, #X11669), Ephrin B3 (santa cruz, # _s_c-514139), PTK7 (Proteintech, #17799), E-cadherin (R&D Systems, #AF748), BCAM (R&D Systems, #BAF148), STAT3 124H6 (cell signaling #9139), p-STAT3 (cell signaling #9131).

### Digestion of heparan sulfate and chondroitin sulfate chains

To enzymatically digest the glycosaminoglycans (GAGs) on SDC, including heparan sulfate (HS) and chondroitin sulfate (CS), nine volumes of cell extract or EVs were incubated with one volume of 10x heparitinase reaction buffer (1M NaCl, 500mM hepes pH 7,0, 10mM CaCl2, 1% TX100, Pepstatin and Leupeptin, both at a concentration of 10 µg/mL), heparitinase (0.4 milliIU, amsbio), and chondroitinase (20milliunits, amsbio). Samples were incubated at 37°C for 4 hours to ensure complete enzymatic digestion. The reaction was terminated by the addition of 5× SDS-PAGE sample buffer. The resulting samples were either processed immediately or stored at −20°C until further analysis by SDS-PAGE and subsequent applications.

### GFP-Trap

MCF7 cells overexpressing GFP-SDC4, or GFP alone as control for 48 h were resuspended in lysis buffer supplemented with 1% detergent (Brij97) for 30 min at 4 °C. Extracts were then centrifuged for 30 min at 10,000 g at 4 °C. Immunoprecipitation was performed for 1 h at 4 °C by incubating GFP-Trap_A beads (Chromtek) with the cellular extracts. After immunoprecipitation, the beads were washed three times in PBS. Proteins coimmunoprecipitated with GFP-SDC4 were detected with corresponding antibodies by Western blot analysis.

### Small extracellular vesicles isolation and total cell lysates

sEVs were collected from equivalent amounts of culture medium, conditioned by equivalent amounts of cells. After the required time of RNAi or drugs treatments, the cell layers were washed twice with PBS and refreshed with DMEM containing 10% EV-downregulated FCS. Cell conditioned media were collected 16 hours later.

#### For differential ultracentrifugation

sEVs were isolated from these media by three sequential centrifugation steps at 4°C: 10 min at 500 x g, to remove cells; 30 min at 10,000 x g, to remove large EVs; and 1.5 hours at 100,000 x g, to pellet small EVs, followed by one wash (suspension in PBS / centrifugation at 100,000 x g). sEV pellets prepared by differential high-speed centrifugation were then re-suspended in PBS. sEVs represent the 100K pellet isolated by differential ultracentrifugation. The corresponding cell layers were washed in cold PBS and scraped on ice in lysis buffer (Tris 30 mM pH 7.4, NaCl 150 mM supplemented with 1% NP40 and protease inhibitor Leupeptin and Pepstatin A (1µg/mL, Sigma Aldrich).

#### For size exclusion chromatography

conditioned media were harvested and centrifuged–at 2,000 g for 20 min at 4°C to discard cell debris in the 2K pellet and then concentrated on a 10 kDa Amicon Filter. Media concentrated to ∼500 μl were overlaid on 35 nm qEVsize-exclusion columns (Izon, qEVoriginal / 35 nm Gen 2) for separation. According to manufacturer’s instructions, SEC fractions were collected in 500 μl volume. For functional experiments fractions F6-F9 were pooled for MCF7 secreting cells or fractions F6-F8 for MDA-MB-468 secreting cells. For western blot analysis, pooled fractions were further concentrated using 10 KDa cut-off filters (Amicon Ultra-4, Millipore).

### Measurement of extracellular vesicles concentration and size

#### Nanoparticle Tracking Analysis (NTA, Nanosight)

Extracellular vesicles were analyzed at similar dilutions with a Nanosight NS-300 instrument (Malvern). 10% of the secretome of approximatively 5 × 10^6^ MCF7 cells were injected for NTA analysis. For each sample, three videos of 60 seconds were recorded at 25 °C and used to calculate mean values of particle concentration.

#### Microfluidic Resistive Pulse Sensing (MRPS, Spectradyne)

Extracellular vesicles concentration was measured using MRPS with the Spectradyne nCS2 following the manufacturer’s instructions. sEV were diluted with PBS in 1% Tween-20 (sample buffer filtered with a 0.02 µm syringe). Samples (5 µL) were placed in microchips (C-400) and inserted into the nCS2. nCS2 Viewer was used for analysis. Briefly, each sample was quantified with dilution factor, background subtracted and saved for graphing purposes. MRPS running buffer (1% Tween-20 added to 1X PBS) was filtered through a 0.2 µm bottle filter.

### Proteomics sample processing and mass spectrometry

Protein samples from Evs analysis or interactome experiments were stacked on NuPAGE™ in a single band to perform trypsin digestion before mass spectrometry analysis using an Orbitrap Fusion Lumos Tribrid Mass Spectrometer (ThermoFisher Scientific, San Jose, CA) in data-independent acquisition mode (DIA) or Data dependent analysis (DDA) respectively. For DIA, Protein identification and quantification was processed using the DIA-NN 1.8 algorithm (Demichev *et al*, 2020) and DIAgui package (https://github.com/marseille-proteomique/DIAgui (Gerault *et al*, 2024)). In case of DDA, proteins identification and quantification were processed using the MaxQuant computational proteomics platform, version 1.6.2.1. The statistical analysis was done with the Perseus program (version 1.6.15.0) (Tyanova & Cox, 2018). The mass spectrometry proteomics data and detailed methods have been deposited to the ProteomeXchange Consortium via the PRIDE partner repository with the data set identifier PXD065942 for interactome experiments or with the data set identifier PXD065960 for the EVs proteome analysis (https://www.ebi.ac.uk/pride/).

### Heat-map generation for mass spectrometry analysis

Protein transmembrane regions and sequences were retrieved using the UniprotR package (v2.4.0). Peptides from each protein were aligned to their corresponding sequences by string matching. The Log2 abundance of peptides was extracted from DIANN reports and is presented in the main part of the heatmap. For each peptide, the KO vs. WT differential abundance was calculated. For a protein, each peptide being located either in the ECD or ICD region, the differential abundances were grouped by region and compared using a t-test. These analyses and visualizations were performed in R 4.4.2 using the stats package (v4.4.2), ComplexHeatmap (v2.22.0) and ggplot2 (v3.5.1). Analysis scripts are available upon request.

### Gene ontology analysis

Gene ontology (GO) enrichment were conducted using Gene Set Enrichment Analysis (GSEA) of WebGestalt (http://www.webgestalt.org/process.php). List of proteins was submitted as a query for GSEA, homo sapiens was selected as the organism and P < 0.05 was used as a selection criterion. The analysis was performed using Reactome Pathway Database (presenting the more abundant functional category supported by WebGestalt 2019 (Liao *et al*, 2019)).

### Endosomal escape assay

Serum-free concentrated media (containing sEVs) of HEK293 or MCF7 cells overexpressing HiBiT-syntenin fusion protein were prepared using Amicon filters and were incubated for 4 hours with recipient MCF7 (plated for 16 h and washed before the start of the experiment with DMEM without FBS) at ratio of 10 producing cells per 1 recipient cell. Recipient cells were washed two or three times, trypsinized, pelleted at 500 x g for 5 mins at 4°C, and split in two halves. One half, to measure the EV uptake was incubated with 2.5 µg of rLgBiT in the presence of 0.8% of Triton X-100 and 8.3 µM/well Furimazine for 5 min before measurement of luminescence. The other half, to measure the cytosolic delivery, was incubated solely with 8.3 µM/well Furimazine before measurement of luminescence. The luminescence was measured on a Victor Nivo plate reader (PerkinElmer).

### Phospho-kinase Array

Proteome Profiler Human Phospho-kinase Array (R&D, ARY003C) was performed according to the manufacturer’s protocol. HUVEC cells were cultured in 10cm dishes in 0,1% gelatin (1 million cells per dish). They were then starved for 3h and then treated for 30min with fresh 100.000 sEVs per target cell (either MCF7/MDA-MB-468 WT sEVs or MCF7/MDA-MB-468 ADAM10-KO sEVs isolated by size exclusion chromatography as previously described). Cell lysates were collected as described in the kit protocol, quantified using PierceTM BCA Protein Assay Kit (Thermo ScientificTM, 23225) and the same amount of protein was loaded on the membrane for each condition, following the manufacturer’s array procedure. Images were captured by ImageQuant800 (Cytiva) and then analyzed using a homemade ImageJ macro, which measured pixel intensity and local background for each spot. The log 2 mean pixel intensity (background subtracted) for each condition were shown in the heatmap. Of note, only proteins showing a change in signal intensity greater than 0 units (after background subtraction), in at least one condition, were included in the analysis.

### Statistical analysis

Differences between groups were determined by using the Kruskal-Wallis one-way non-parametric ANOVA test. Multiple comparisons were carried out using the two-way ANOVA test, unpaired two-tailed t-test or multiple unpaired t-test. Single comparisons were carried out using the non-parametric unpaired Mann–Whitney test. * P < 0.05, ** P < 0.01, *** P < 0.001, **** P < 0.0001, n.s. non-significant. All statistical analyses were conducted using GraphPad Prism v10.0c software.

## Acknowledgments

We thank Dr. Cyril Addi for the GFP-SDC4 construct, Dr. Eric Rubinstein for ADAM10 and tetraspanin antibodies and Dr. Jean-Hugues Guervilly for their scientific advice.

This work was supported by grants from La Ligue Nationale Contre le Cancer (EL2018.LNCC/PaZ and EL2024.LNCC/PaZ), the French National Research Agency (ANR-18-CE13-0017, Project SynTEV and ANR-24-CE13-1161-01), the Canceropole Provence-Alpes-Cote-d’Azur (2024-06Kpole) and KU Leuven Internal Funding (C14/20/105 and C14/25/141). LH is the recipient of an FWO aspirant fellowship number 1SE2422N.

We are grateful to the extracellular vesicle core facility of the CRCM for providing services and equipments (CEEVEC, Centre d’Expertises et d’Expérimentations sur les Vésicules Extracellulaires en Cancérologie).

Proteomics analyses were done using the mass spectrometry facility of Marseille Proteomics (marseille-proteomique.univ-amu.fr) supported by IBISA (Infrastructures Biologie Santé et Agronomie), Plateforme Technologique Aix-Marseille, the Cancéropôle PACA, the Provence-Alpes-Côte d’Azur Region, the Institut Paoli-Calmettes, Fonds Européen de Développement Regional (FEDER) and Plan Cancer. Some figures were created with BioRender.com.

## Notes

### Competing Interest Statement

The authors have declared no competing interest.

### Summary of Updates

After submission, I realized that I made an error in the author order entered in the submission system. Specifically, Pascale Zimmermann should be listed as the last author on the bioRxiv website, as correctly indicated in the uploaded PDF version of the manuscript.

